# AtNHR2A and AtNHR2B participate in unconventional protein secretion in response to environmental stress

**DOI:** 10.64898/2026.09.10.750635

**Authors:** Ha Thi Kim Nguyen, Thiago Maia, Biwesh Ojha, Clemencia M. Rojas

## Abstract

The *Arabidopsis thaliana* <u>n</u>on<u>h</u>ost resistance proteins 2A (AtNHR2A) and 2B (AtNHR2B) play crucial roles in plant immunity as the single mutants *Atnhr2a* and *Atnhr2b* and the double mutant *Atnhr2bAtnhr2a* are susceptible to the non-adapted pathogen *Pseudomonas syringae* pv. tabaci that is unable to infect wild-type Col-0 plants. The localization of fluorescent versions of AtNHR2A and AtNHR2B to compartments of the endomembrane system together with their interaction with secreted proteins suggested a function in endomembrane-mediated secretory processes participating in plant immunity. Comparative apoplastic proteomics analysis between wild type Col-0 and the double mutant *Atnhr2bAtnhr2a* after treatment *P. syringae* pv. tabaci, revealed that AtNHR2A and AtNHR2B are indeed required for the secretion of proteins containing N-terminal signal peptides that occurs through the conventional protein secretion pathway. In this work, we leveraged these apoplastic proteomics datasets to identify proteins lacking N-terminal signal peptide and expected to be secreted through unconventional secretion pathway(s). We discovered that AtNHR2A and AtNHR2B are also required for the secretion of proteins through an unconventional secretion pathway that, intriguingly, included proteins previously associated with abiotic stress. These findings led us to define the subcellular dynamics of AtNHR2A and AtNHR2B, and through co-localization analyses and the use of vesicle trafficking inhibitors, we uncovered their trafficking pathways transitioning through Golgi-dependent and Golgi-independent pathways to ultimately reach the central vacuole. Our findings suggest that AtNHR2A and AtNHR2B participate in a multivesicular bodies-vacuole-mediated unconventional secretion pathway that results in the release of proteins involved in plant responses to environmental stresses.

## Introduction

Plants respond to infectious agents through complex signal transduction pathways that are activated when plants recognize a potential pathogen. Pathogen recognition occurs in the extracellular milieu through Pathogen Recognition Receptors (PRR) that detect conserved features in pathogens called Pathogen Associated Molecular Patterns (PAMPs), or intracellularly by Nucleotide-binding and Leucine-rich-repeat-containing receptors (NLRs) that detect pathogen proteins called effectors, triggering PAMP-triggered immunity (PTI) and Effector-trigger immunity (ETI), respectively (Jones et al., 2024, Jones and Dangl, 2006). PTI and ETI synergistically trigger a plethora of events aimed at controlling pathogen proliferation and minimizing cellular damage (Ngou et al., 2021, Yuan et al., 2021). These events include: 1) the remodeling of the cell wall, presumably to provide strength and to prevent nutrient leakage to the apoplast, and 2) the release to the apoplast of powerful cocktails of defense-related proteins with antimicrobial properties (Bhandari and Brandizzi, 2024, Del Corpo et al., 2024). Both events have been recognized for many years and are known to occur through secretory pathways of the endomembrane system comprising several membrane-bound interconnected organelles including endoplasmic reticulum (ER), Golgi apparatus, trans-Golgi network (TGN), multivesicular bodies (MVBs) and vacuole (Bhandari and Brandizzi, 2024, Gu et al., 2017, Kwon et al., 2008).

Two main protein secretion pathways are known: conventional protein secretion (CPS) pathway, and unconventional protein secretion (UPS) pathway. In plants, as well as in other eukaryotes, proteins destined to the CPS are synthesized with an N-terminal secretion signal and packaged into vesicles as cargo. Such vesicles are then translocated sequentially from the ER to Golgi apparatus and TGN through membrane-fusion events. Ultimately, fusion of transporting vesicles with the plasma membrane (PM) releases cargo proteins to the apoplastic space (Davis et al., 2016, Goring and Di Sansebastiano, 2018). In contrast, proteins secreted through UPS are synthesized without a signal peptide (leaderless), and either directly reach the PM to be released to the apoplast, or become cargo in vesicles that bypass Golgi apparatus and fuse with other organelles such as MVB, vacuole, and exocyst-positive organelle (EXPO); similar to CPS, fusion of these organelles with the PM releases cargo proteins to the apoplast (Davis et al., 2016, Krause et al., 2013, Ding et al., 2014, Wang et al., 2018). Interestingly, the finding that leaderless proteins are secreted in higher abundance in conditions of biotic and abiotic stress in comparison with non-stress conditions, highlights the importance of the UPS pathway mediating cellular responses to environmental stress (Agrawal et al., 2010). Despite its importance, the function of the UPS pathway in stress responses is still not fully understood, nor whether specific subcellular compartments are associated with responses to specific types of stress.

We previously identified two Arabidopsis proteins, AtNHR2A (*Arabidopsis thaliana* nonhost resistance 2A) and AtNHR2B (*Arabidopsis thaliana* nonhost resistance 2B), as immune-related proteins localized to compartments of the endomembrane system (Singh et al., 2018). We further showed that AtNHR2A and AtNHR2B interact with proteins known to be secreted (Singh et al., 2020), and that both AtNHR2A and AtNHR2B are required for the secretion of defense-related proteins to the apoplast through the CPS pathway in response to inoculation with the non-adapted pathogen *Pseudomonas syringae* pv. tabaci (*Pstab*) (Maia et al., 2026).

In this study, we found that AtNHR2A and AtNHR2B are also required for protein secretion through the UPS pathway for the apoplastic delivery of proteins associated with plant responses to both biotic and abiotic stresses. Moreover, through transient co-localization and quantitative live-cell imaging assays in *Nicotiana benthamiana,* we defined that both proteins, AtNHR2A and AtNHR2B, transition through Golgi-dependent and Golgi-independent trafficking pathways, in agreement with their function in CPS and UPS, and uncovered a MVB-vacuole-mediated UPS pathway for these proteins. Collectively, the data highlights the function of AtNHR2A and AtNHR2B orchestrating protein trafficking to MVB and vacuole that precedes secretion to the apoplast of proteins associated with plant responses to environmental stresses.

## Materials and methods

### Bacterial strains and plasmids

*Agrobacterium tumefaciens* strains GV3101, GV2260 and C58C1 harboring plasmids of interest used in this study are listed in Supplementary Table S1. *A. tumefaciens* strains were grown in Luria-Bertani (LB) medium supplemented with appropriate antibiotics, including Rifampicin (25 µg/mL), Spectinomycin (50 µg/mL) and Kanamycin (50 µg/mL).

Construct *GONST1-GFP* driven by the *35S* promoter was generated by amplifying *GONST1* from plasmid *mEOS-GONST1* (Mathur et al., 2010) by PCR, using forward primer, gggACAAGTTTGTACAAAAAAGCAGGCTggatgaaattgtacgaacacgatg, and reverse primer, ggggACCACTTTGTACAAGAAAGCTGGGTgggacttctccctcattttgg. PCR product was cloned into *pDONR-Zeo* by Gateway BP clonase^TM^reaction and then transferred into *pMDC83* via Gateway LR clonase^TM^ reaction.

Constructs *γTIP-mCherry* and *AtPIP2A-mCherry* were obtained from Arabidopsis Biological Resource Center (ABRC, The Ohio State University).

### Transient expression in *Nicotiana benthamiana* and live cell imaging

*Nicotiana benthamiana* plants were grown under long-day condition (16/8 h, light/dark) at 25°C, and 50% humidity.

*Agrobacterium* strains harboring constructs of interest were grown at 28°C and 200 rpm in a shaker incubator. Overnight cultures were harvested by centrifugation at 6,000 rpm, bacterial pellets were re-suspended in induction buffer (30mM MES, 0.25% mannitol, 200µM acetosyringone) and incubated for 3 h at 28°C with constant aeration. Induced cultures were harvested by centrifugation at 6,000 rpm, resuspended in 10mM MES buffer (pH 5.5) and bacterial concentration adjusted to an OD_600_= 0.3. *Agrobacteria* strains were used to infiltrate the abaxial side of three-week-old *N. benthamiana* plants. For treatment with vesicle trafficking inhibitors, *N. benthamiana* leaves expressing proteins of interest were infiltrated with Brefeldin A (BFA, Sigma-Aldrich, 50 µg/mL) and imaged at 1h and 3h after infiltration (hai), or Wortmannin (WM, Sigma-Aldrich, 25 µM) and imaged at 30 min after infiltration. Infiltrated leaves were imaged at two to three days after infiltration on a Nikon A1 upright confocal microscope using 60X magnification and collecting images in the GFP (excitation: 488 nm, emission: 507nm), or RFP/mCherry (excitation: 587 nm, emission: 610nm) channels.

### Image quantification

Image quantification was conducted by Image J. To assess colocalization, raw images were imported into ImageJ and the green and red fluorescence channels were merged. Colocalization scores were measured using Coloc2 method with Costes algorithm automatic threshold, and collecting Pearson’s Correlation Coefficient (PCC), Thresholded Manders’ Split Coefficients tM1 and tM2 with Costes P-value. Only image pairs with a Costes P-value ≥ 0.95 were retained, confirming that the observed correlation was significantly greater than that of randomized images. Coefficient values were interpreted according to the following ranges: anti correlation (PCC= -1 to -0.1), no co-localization (PCC = -0.1 to 0.1, tM= 0 to < 0.2), weak colocalization (PCC= +0.1 to < +0.3, tM= 0.2 to <0.5), moderate colocalization (PCC=+0.3 to <+0.5, tM= 0.5 to <0.8) and strong colocalization (PCC= +0.5 to +1, tM=0.8 to 1) (Barlow et al., 2010, Zinchuk et al., 2013). When PCC and tM fell within the same category, colocalization was scored at that level; when they fell into different categories, the lower of the two was reported, providing a conservative estimate.

## Results

### AtNHR2A and AtNHR2B contribute to protein secretion via unconventional protein secretion pathway

Our previous comparative apoplast proteomics analysis between wild-type Col-0 and the double mutant *Atnhr2bAtnhr2a* after inoculation with *Pstab* identified 2,651 proteins in the apoplastic fluids, but only 8.1% of them contained an N-terminal signal peptide (Maia et al., 2026), suggesting that a significant number of apoplastic proteins may be secreted through the UPS pathway. To identify candidate UPS proteins and further evaluate whether their secretion is dependent on AtNHR2A and AtNHR2B function we employed a comparative proteomics approach using our apoplastic and whole leaf proteome dataset. First, we retrieved the average normalized intensity-Based Absolute Quantification (iBAQ) values of the secreted proteins with N-terminal signal peptides previously identified in the apoplastic and whole leaf proteome datasets obtained after inoculating Arabidopsis wild-type Col-0 and the double mutant *Atnhr2bAtnhr2a* with *Pstab* (Figure 1A) and then calculated the average iBAQ fold change (apoplast/whole leaf) for proteins detected in each genotype/time point combination. We found that those fold changes were 27-fold for Col-0 at 24 hai, 39-fold for Col0 at 72 hai, 35-fold for *Atnhr2bAtnhr2a* at 24 hai, and 34-fold for *Atnhr2bAtnhr2a* at 72 hai (Figure 1B). We used the lowest enrichment observed (27-fold) to establish an apoplast/whole-leaf enrichment cut-off of 25. Therefore, proteins lacking a predicted N-terminal signal peptide and exhibiting apoplast/whole-leaf iBAQ ratios greater than 25 were considered candidate UPS proteins.

**Figure 1.**
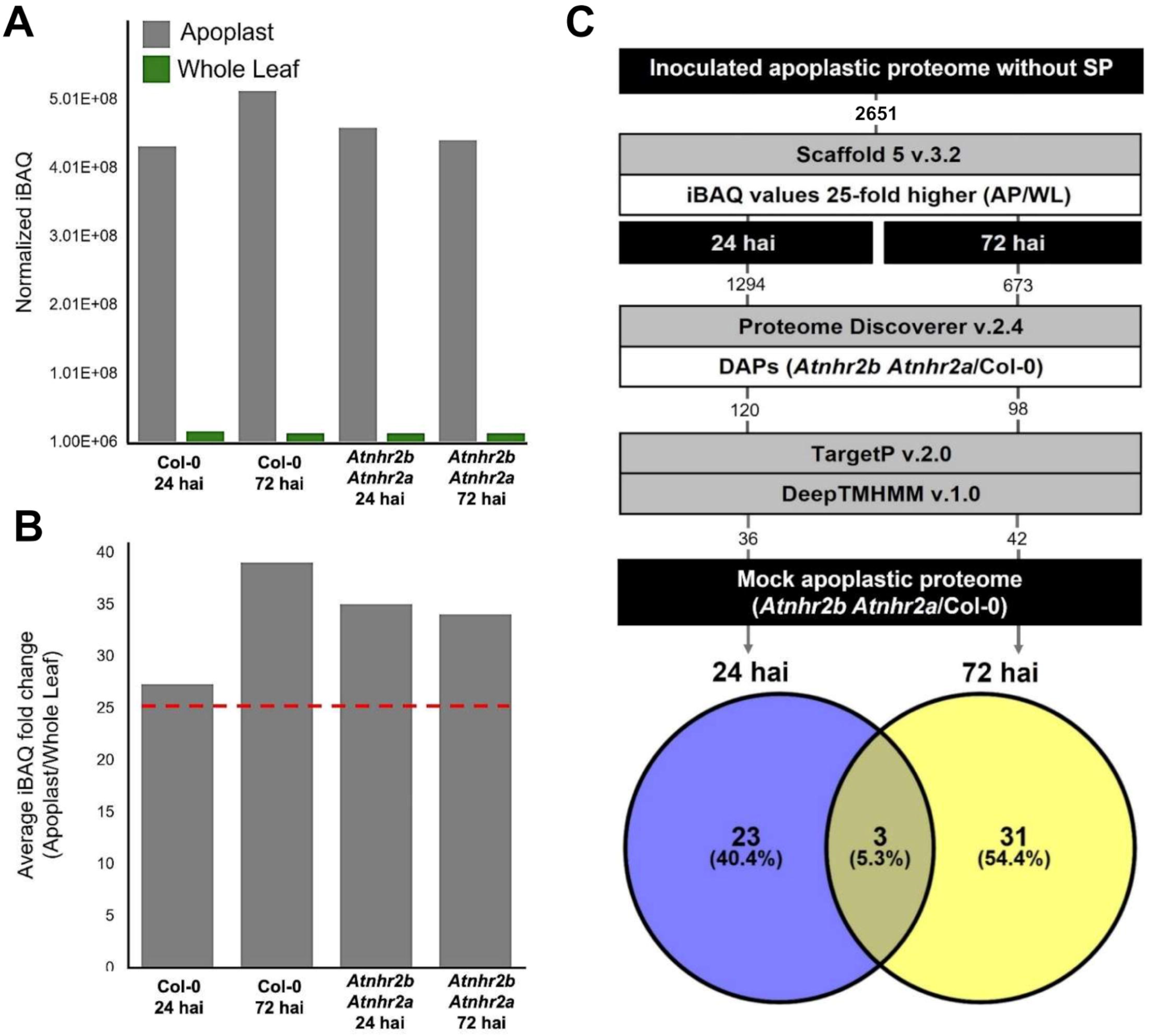
Identification of Differential Abundance Proteins (DAPs) lacking N-terminus signal peptides. Average normalized iBAQ values of proteins containing N-terminal signal peptides in both apoplast (grey bars) and whole leaf (green bars) proteome datasets (A) were used to identify proteins lacking N-terminal signal peptide based on their respective Apoplast/whole leaf ratios (B). Workflow for the identification of Differential Abundance Proteins (DAPs) without signal peptides in the apoplast of *Atnh2b Atnh2a* compared to Col-0 after inoculation with *Pseudomonas syringae* pv. tabaci (Pstab at 24 and 72 hours after inoculation (hai) (C).

After calculating the average apoplast/whole leaf ratios using iBAQ value for those 2,651 proteins, we selected 1,294 proteins at 24 hai and 673 proteins at 72 hai that exhibited apoplast/whole leaf ratios at least greater than 25 (Figure 1C). Next, we analyzed these protein datasets to identify Differential Abundance Proteins (DAPs) in the apoplast between *Atnhr2bAtnhr2a* and Col-0 and found 120 DAPs at 24 hai and 98 DAPs at 72 hai. We also examined at the abundance ratio (*Atnhr2bAtnhr2a* vs. Col-0) of these DAPs in the mock-treated apoplastic proteome to select only those DAPs with log2 fold changes that differed by >1 (for proteins with increased abundance) or <-1 (for proteins with decreased abundance) between the *Pstab*-inoculated and mock-treated samples in the final dataset. This filtering process led to the identification of 26 DAPs at 24 hai and 34 DAPs at 72 hai, with 3 DAPs common to both time points (Figure 1C, Supplementary Tables S2 and S3).

Hierarchical cluster analysis of the 26 DAPs lacking signal peptides identified at 24 hai, highlighted distinct patterns of protein abundance across the treatments and revealed six main clusters (Figure 2A, Supplementary Table S2). Cluster 1 includes five DAPs, that showed higher abundance in *Atnhr2bAtnhr2a* mutant under mock treatment in comparison with Col-0, but four of those DAPs showed reduced abundance in the *Atnhr2bAtnhr2a* mutant in comparison with Col-0 after inoculation with *Pstab* (Figure 2A, Supplementary Table S2). Cluster 2 includes three DAPs that showed higher abundance in Col-0 than in *Atnhr2bAtnhr2a* under mock treatment, but only two showed reduced abundance in the *Atnhr2bAtnhr2a* mutant after inoculation with *Pstab* (Figure 2A, Supplementary Table S2). Remarkably clusters 3, 4 and 5, collectively included 15 DAPs that have a mix combination of abundance between Col-0 and *Atnhr2bAtnhr2a* under mock treatment, but all those DAPs showed significantly lower abundance in the *Atnhr2bAtnhr2a* when compared with Col-0 after *Pstab* inoculation (Figure 2A, Supplementary Table S2). Cluster 6 includes three DAPs that showed higher abundance in *Atnhr2bAtnhr2a* in comparison with Col-0 under both mock treatment and *Pstab* inoculation (Figure 2A, Supplementary Table S2). Altogether, we identified 21 DAPs with reduced abundance in *Atnhr2bAtnhr2a* compared to Col-0 after *Pstab* inoculation. We classified these DAPs into four functional categories based on Gene Ontology analysis and our manual curation of the published literature. Those categories are: 1) Responses to abiotic stress (8 DAPs), 2) Responses to biotic stress (3 DAPs), 3) Responses to both biotic and abiotic stress (2 DAPs) and 4) Plant Development (2 DAPs). Six additional DAPs with predicted functions but not validated in the literature, were classified as unknown. Notably, among the stress-responsive DAPs, 7 were also responsive to the stress hormone abscisic acid (ABA) (Table 1).

**Figure 2.**
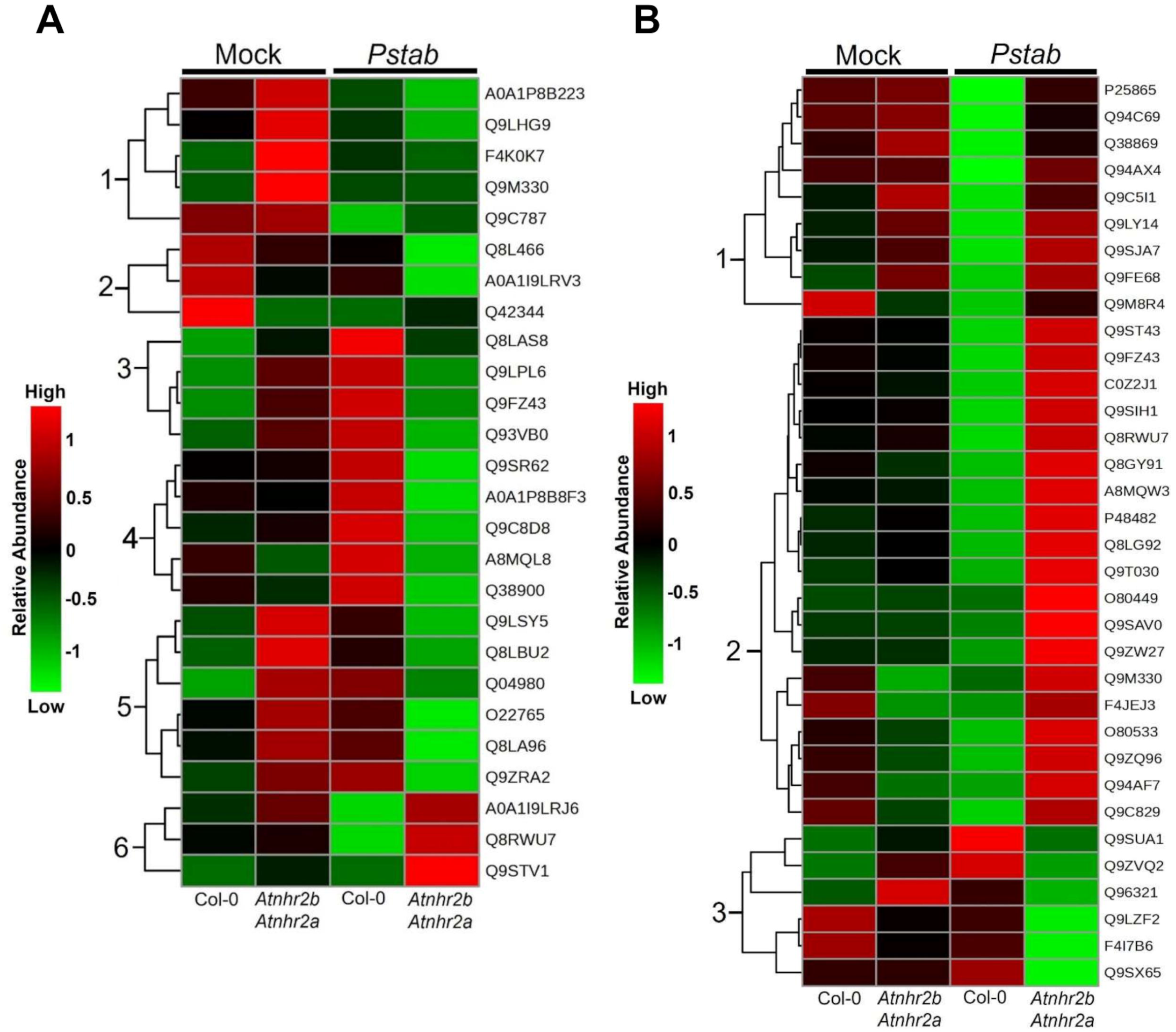
Hierarchical clustering analysis of the DAPs without N-terminal signal peptides after *Pseudomonas syringae* pv. tabaci inoculation. Hierarchical clustering analysis of the 26 DAPs and 34 DAPs without N-terminal signal peptides identified 24 hours (A) and 72 hours (B), respectively, in Col-0 and *Atnhr2bAtnhr2a* under mock-treatment and *Pseudomonas syringae* pv tabaci inoculation. Color scale represents relative abundance of each DAP.

**Table 1.** Differential Abundance Proteins (DAPs) without N-terminal signal peptides with lower abundance in the apoplast of *Atnhr2bAtnhr2a* compared to Col-0 at 24 and 72 h after inoculation with *Pseudomonas syringae* pv. tabaci.

| Gene ID/ Protein Accession | Description | Abundance ratio* | Biological activity** | Reference*** |
| --- | --- | --- | --- | --- |
| **A) DAPs identified at 24h after inoculation with *P. syringae* pv. *tabaci*.** |  |  |  |  |
| **1) Response to abiotic stress** |  |  |  |  |
| At5g66400<br>F4K0K7 | Responsive to ABA18<br>(RAB18) | -6.6 | Response to drought and cold stresses. Response to ABA. | Lång & Palva, 1992, Lang et al., 1994, Jeannette et al., 1999. |
| At5g52300<br>Q04980 | Responsive to desiccation<br>29B (RD29B) | -2.3 | Response to cold, salt, and desiccation. Response to ABA. | Yamaguchi-Shinozaki & Shinozaki, 1993, Nordin et al., 1993, Msanne et al., 2011, Kaur et al., 2025 |
| At4g36020<br>A0A1P8B8F3 | Cold shock domain protein 1<br>(CSDP1) | -6.6 | Response to cold and salt. | Yang & Karlson, 2013, Park et al., 2009, Juntawong et al., 2013 |
| At5g54080<br>Q9ZRA2 | Homogentisate 1,2-dioxygenase (HGO) | -6.6 | Response to salt stress. | Han et al., 2013, Huang et al., 2018 |
| At3g21790<br>Q9LSY5 | UDP-glycosyltransferase<br>71B7 (UGT71B7) | -6.6 | Response to dehydration, osmotic and high-salinity stresses. Control endogenous ABA level in plant cells. | Dong et al., 2014 |
| At5g15230<br>A8MQL8 | GA-stimulated Arabidopsis 4<br>(GASA4) | -1.9 | Exhibits redox activity. Response to heat stress, light stress, ABA. | Ko et al., 2007, Rubinovich & Weiss, 2010, Chen et al., 2007, Qu et al., 2016 |
| At1g21380<br>Q9LPL6 | TOM1-like protein 3 (TOL3) | -6.6 | Response to drought stress. Reduction of stomatal apertures. Induced by ABA. | Korbei et al., 2013, Moulinier-Anzola et al., 2014 |
| At1g63460<br>Q8LBU2 | Probable glutathione<br>peroxidase 8 (GPX8) | -6.6 | Response to oxidative, heat, high light stress. | Gaber et al., 2012, Gaber, 2014, Passaia et al., 2014 |
| **2) Response to biotic stress** |  |  |  |  |
| At3g53970<br>Q9M330 | Proteasome regulator 1<br>(PTRE1) | -6.6 | Regulator of plant immunity. Response to auxin, ABA. | Yang et al., 2016, Thulasi Devendrakumar et al., 2019, Hao et al., 2026 |
| At3g46000<br>A0A1I9LRV3 | Actin depolymerizing factor 2<br>(ADF2) | -1.9 | Response to nematode infection. Control actin depolymerization. | Ruzicka et al., 2007, Clement et al., 2009 |
| At2g41530<br>Q8LAS8 | S-formylglutathione hydrolase (SFGH) | -1.3 | RNA is upregulated in root of nematode infection. Formaldehyde catabolic process for detoxification. | Kordic et al., 2002, Hütten et al., 2015 |
| **3) Response to both abiotic stress and biotic stress** |  |  |  |  |
| At2g16600<br>Q38900 | Peptidyl-prolyl cis-trans isomerase (ROC3) | -1.3 | Response to drought stress and ABA. Response to *P. syringe*. | Liu et al., 2021, Pogorelko et al., 2014, Luo et al., 2024 |
| At4g17720<br>Q8LA96 | BPA1-like 1 (PBL1) | -2.9 | Response to salt and glucose and biotic stress. | Li et al., 2019, Palm et al., 2019 |
| **4) Plant development** |  |  |  |  |
| At3g09970<br>Q9SR62 | Rhizobiale-like phosphatase 2 (RLPH2) | -6.6 | A regulator of seed dormancy. Involved in ABA and GA pathways. | Labandera et al., 2026, Uhrig et al., 2016 |
| At4g02610<br>O22765 | Tryptophan synthase alpha chain (TSAα) | -6.6 | Auxin biosynthesis. | Radwanski et al., 1995 |
| **Unknown function** |  |  |  |  |
| At1g24050<br>Q8L466 | Lsm domain-containing protein | -6.6 | RNA processing |  |
| At1g66070<br>Q9C8D8 | Eukaryotic translation initiation factor 3 | -6.6 | Translation initiation factor |  |
| At2g40660<br>Q93VB0 | Nucleic acid-binding, OB-fold-like protein | -6.6 | Aminoacyl-tRNA synthetase |  |
| At2g35820<br>A0A1P8B223 | Ureidoglycolate hydrolase | -6.6 | Hydrolase |  |
| At3g12390<br>Q9LHG9 | Nascent polypeptide-associated complex | -2 | Protein transport |  |
| At1g16810<br>Q9FZ43 | 7-dehydrocholesterol reductase-like protein | -6.6 | Others |  |

| **B) DAPs identified at 72h after inoculation with *P. syringae* pv. *tabaci*.** |  |  |  |  |
| --- | --- | --- | --- | --- |
| At4g26630<br>Q9SUA1 | DEK domain-containing<br>protein 3 (DEK3) | -6.6 | Response to salt stress,<br>oxidative stress. ABA-dependent protein | Waidmann et al., 2014, Waidmann<br>et al., 2022. |
| At1g10940 F4I7B6 | Protein kinase (SnRK2.4) | -1.5 |  | Kulik et al., 2012, Mazur et al.,<br>2021, McLoughlin et al., 2012,<br>Adamo et al., 2025 |
| At2g02400<br>Q9ZVQ2 | NAD(P)-binding Rossmann-fold superfamily protein<br>(CCR-like 1) | -6.6 | Putative lignin biosynthesis | Raes et al., 2003 |
| At1g48320<br>Q9SX65 | 1,4-dihydroxy-2-naphthoyl-CoA thioesterase 1<br>(DHNAT1) | -6.6 | Phylloquinone (Vitamin K1)<br>biosynthesis | Widhalm et al., 2012, Furt et al.,<br>2013 |
| At3g06720Q96321 | Importin subunit alpha-1<br>(IMPA-1) | -6.6 | Protein import into nucleus | Hübner et al., 1999 |
| At5g03370<br>Q9LZF2 | Acylphosphatase | -3.1 | Unknown function |  |

\*Abundance ratio Atnht2bAtnhr2a/Col-0 values ( $p$ -value < 0.05)
\*\*Gene ontology annotation and references
\*\*\* Refence list was provided in Supplementary reference

At 72 hai, three main clusters were observed (Figure 2B). Cluster 1 includes nine DAPs with similar abundance levels between Col-0 and *Atnhr2bAtnhr2a* under mock treatment but have higher abundance in the *Atnhr2bAtnhr2a* mutant in comparison with Col-0 after *Pstab* inoculation (Figure 2B, Supplementary Table S3). Cluster 2 includes 19 DAPs that in general, showed similar abundance between Col-0 and *Atnhr2bAtnhr2a* under mock treatment, and like cluster 1 also have higher abundance in *Atnhr2bAtnhr2a* in comparison with Col-0 after *Pstab* inoculation (Figure 2B, Supplementary Table S3). Cluster 3 contains six DAPs, three of them showing higher abundance in Col-0 versus *Atnhr2bAtnhr2a* and three of them showing lower abundance between Col-0 and *Atnhr2bAtnhr2a* under mock treatment; however, after *Pstab* inoculation all these DAPs exhibited lower levels in the *Atnhr2bAtnhr2a* mutant in comparison with Col-0 after (Figure 2B, Supplementary Table S3). Those six proteins in cluster 3 represent proteins secreted in an AtNHR2A- and AtNHR2B-dependent manner. Functional annotations, combined with a survey of the published literature on these DAPs indicated that two of them, DEK domain-containing protein 3 (DEK3) and Protein kinase (SnRK2.4) are associated with abiotic stress, particularly salt stress; two other DAPs encoding NAD(P)-binding Rossmann-fold superfamily protein (CCR-like 1) and 1,4-dihydroxy-2-naphthoyl-CoA thioesterase 1 (DHNAT1) participate in biosynthesis of lignin and Phylloquinone, respectively; one DAP encoding Importin subunit alpha-1 (IMPA-1), implicated in protein import to nucleus and, one DAP predicted to encode an acylphosphatase that remain to be characterized (Table 1).

To investigate whether the differences in abundance of proteins without signal peptides in the apoplast between *Atnhr2bAtnh2a* and Col-0 were due to impairments in the unconventional secretion pathway rather than protein synthesis, we compared the abundance of selected UPS DAPs between apoplast and whole leaf proteomes after *Pstab* inoculation. For this analysis, we selected 15 DAPs with reduced abundance in the apoplast of the double mutant *Atnhr2bAtnhr2a* (Table 1) and that were also detected in the whole leaf proteome (Supplementary Table S4). Among these 15 DAPs, only two DAPs, Homogentisate 1,2-dioxygenase (Q9ZRA2) and importin subunit alpha-1 (Q96321) exhibited significantly lower abundance (*p*-value < 0.05) in the whole leaf samples of the *Atnhr2bAtnhr2a* mutant when compared to Col-0 (Figure 3), the additional 13 DAPs showed reduced abundance in the apoplast of *Atnhr2bAtnhr2a* when compared to Col-0 but exhibited higher or similar abundance (no significant change) in whole leaf samples when comparing the *Atnhr2bAtnh2a* mutant with Col-0, indicating that there is no difference in those protein levels between Col-0 and *Atnhr2bAtnhr2a* (Figure 3). These results indicate that the majority (87%) of UPS DAPs accumulated in the apoplastic compartment are influenced by the absence of functional AtNHR2A and AtNHR2B.

**Figure 3.**
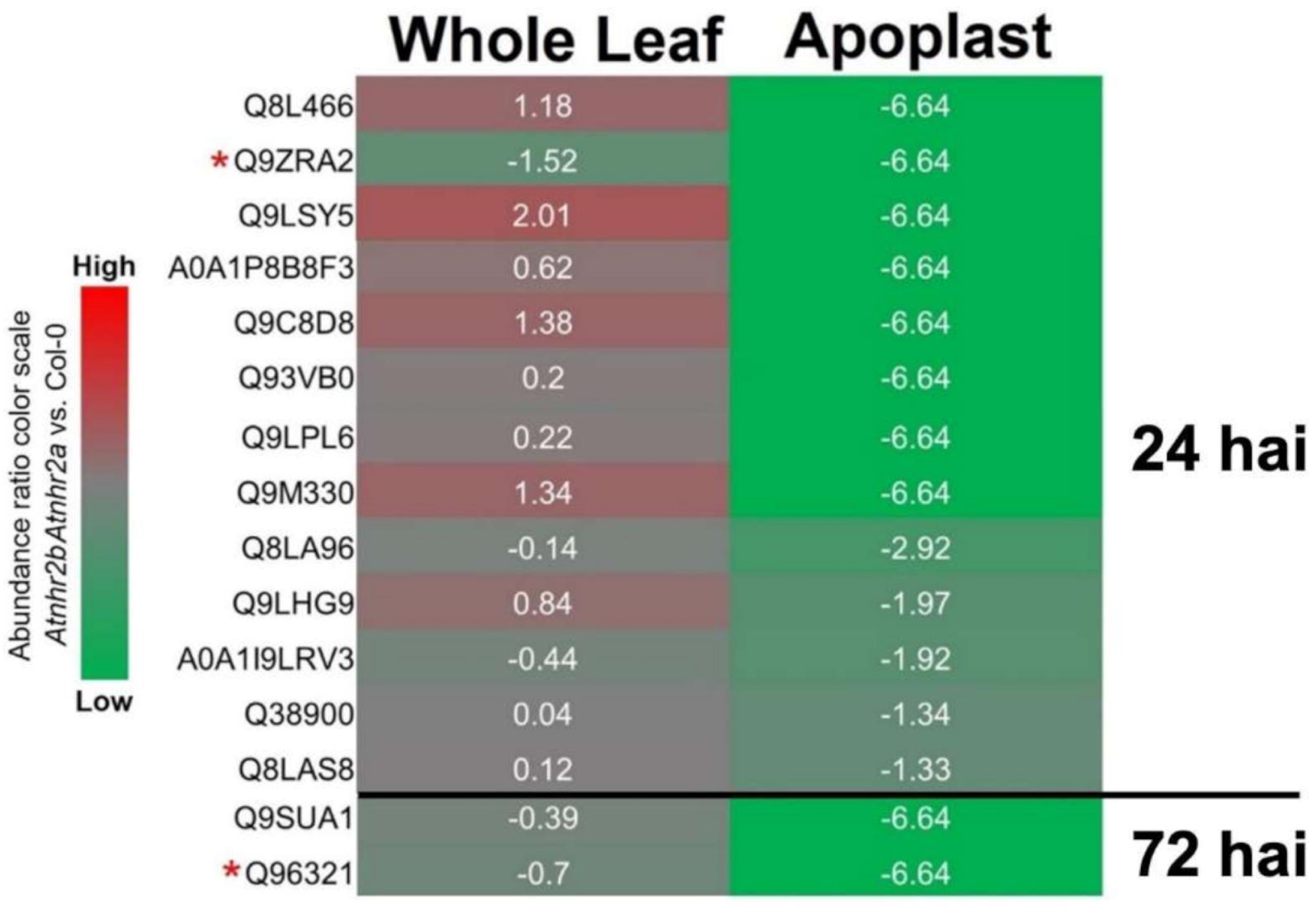
Comparison of abundance ratios of DAPs without signal peptide between *Atnhr2bAtnhr2a* versus Col-0 in whole leaf and apoplast proteomes. Comparison of abundance ratios (*Atnhr2bAtnhr2a* versus Col-0) of DAPs lacking signal peptides identified in both the apoplast and the whole leaf proteomes at 24 and 72 hours after *Pseudomonas syringae* pv. tabaci inoculation. The color scale indicates abundance ratios. Red asterisks denote DAPs with statistically lower abundance (*p*-value < 0.05) in the whole leaf proteome.

### AtNHR2A and AtNHR2B transit through both Golgi-dependent and Golgi-independent routes

Our findings that AtNHR2A and AtNHR2B participate in both CPS and UPS prompted our interest to dissect the trafficking pathways associated with AtNHR2A and AtNHR2B function. To investigate this function, we sought to define their specific subcellular localization by co-expressing fluorescent versions of AtNHR2A and AtNHR2B with fluorescent cellular markers labeling ER (AtWAK2, *Arabidopsis thaliana* wall-associated kinase 2) (Nelson et al., 2007) and Golgi (GONST1, Golgi nucleotide sugar transporter 1) (Baldwin et al., 2001), compartments associated with the early stages of the secretory pathway. Co-expression of *AtNHR2A-GFP* and *AtNHR2B-GFP* with the ER marker *AtWAK2* fused to mCherry revealed co-localization as shown by the overlap of green and red signals. This colocalization is considered moderate based on Pearson’s Correlation Coefficient (PCC) scores of 0.45±0.09 and 0.47±0.12, respectively, and tM1 and tM2 values ranging between 0.6-0.8, indicating moderate localization of AtNHR2A and AtNHR2B to the ER (Figure 4A, Supplementary Table S5). Co-expression of *AtNHR2A-RFP* and *AtNHR2B-RFP* with the Golgi marker, *GONST1* fused with *GFP*, revealed low percentage of co-localization with PCC scores, of 0.28±0.04 and 0.31±0.06 for AtNHR2A and AtNHR2B, respectively, as well as tM1 and tM2 value ranging between 0.5-0.6 (Figure 4B, Supplementary Table S5), indicating partial localization of AtNHR2A and AtNHR2B to Golgi.

**Figure 4.**
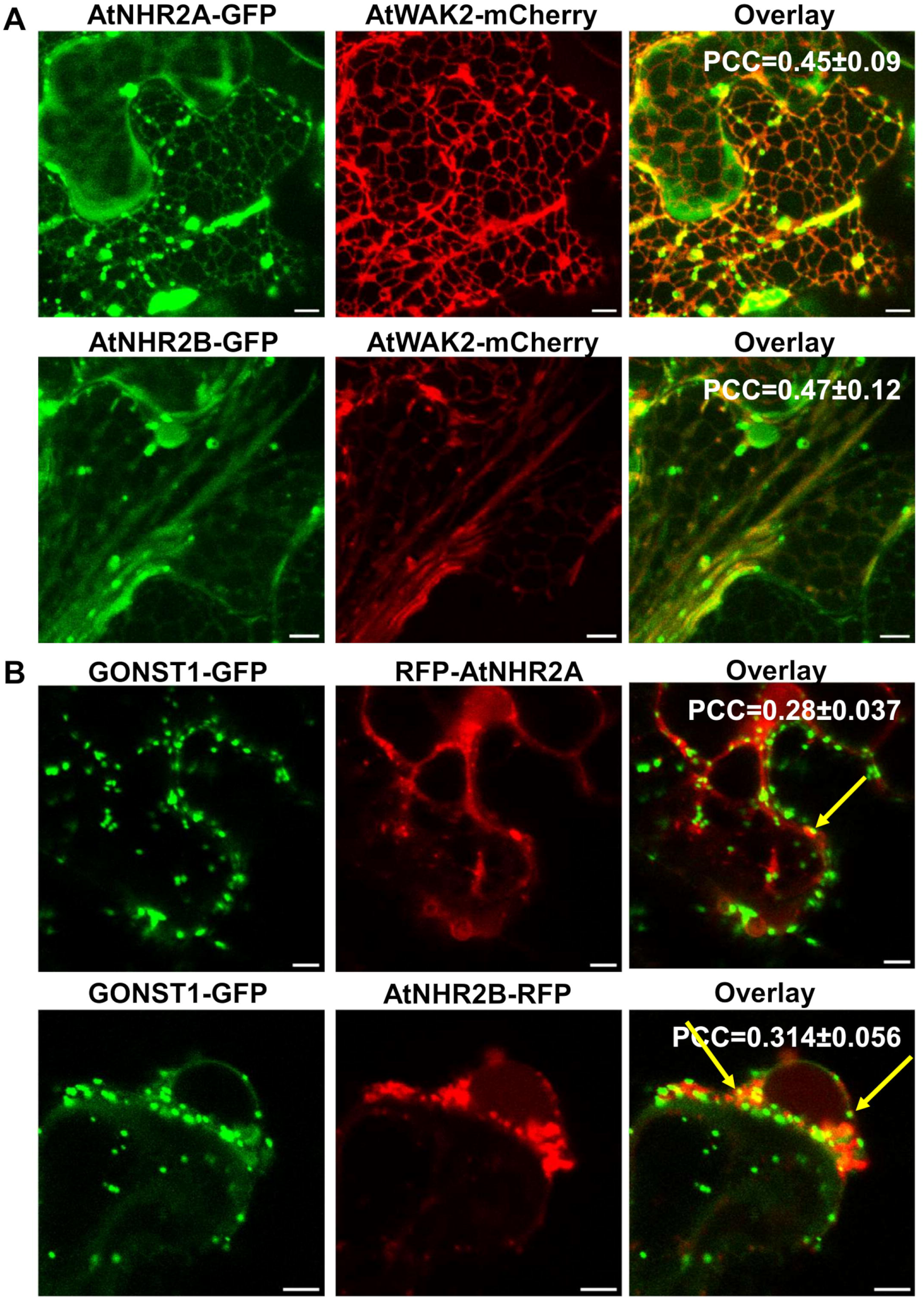
AtNHR2A and AtNHR2B are partially localized to ER and Golgi. Transient co-expression in *Nicotiana benthamiana* was used to evaluate co-localization of fluorescent versions of AtNHR2A AtNHR2B with the ER (A) and Golgi (B), using fluorescent markers AtWAK2-mCherry and GONST1-GFP, respectively. Infiltrated leaves were imaged 2 days after infiltration on a laser scanning confocal microscopy in the green and red channels. Pictures are representative images. Pearson’s correlation coefficient (PCC) value represents average of 10 microscopic fields. Scale bar = 5µm. Images in (A) are Z-stacks of 10 sections. Images in (B) are single scanning.

To confirm the transition of AtNHR2A and AtNHR2B from ER to Golgi, *N. benthamiana* leaves transiently expressing *AtNHR2A-GFP* and *AtNHR2B-GFP* were treated with the vesicle trafficking inhibitor Brefeldin A (BFA) that interferes with the movement of proteins from the ER to Golgi (Ritzenthaler et al., 2002). The results showed the tobacco leaves expressing *GONST1-GFP* followed by infiltration with 0.1% DMSO control resulted in the typical localization of *GONST1-GFP* to Golgi (Figure 5, blue arrow). Transient expression of *AtNHR2A-GFP* and *AtNHR2B-GFP* followed by mock treatment with 0.1% DMSO treatment also resulted in fluorescent signal accumulation in Golgi (Figure 5, blue arrows) as well as in the small punctae previously described (Singh et al., 2018, Marín-Ponce et al., 2023) (Figure 5, white arrows). The ER localization of AtNHR2B was also observed under 0.1% DMSO treatment (Figure 5, yellow arrow).

**Figure 5.**
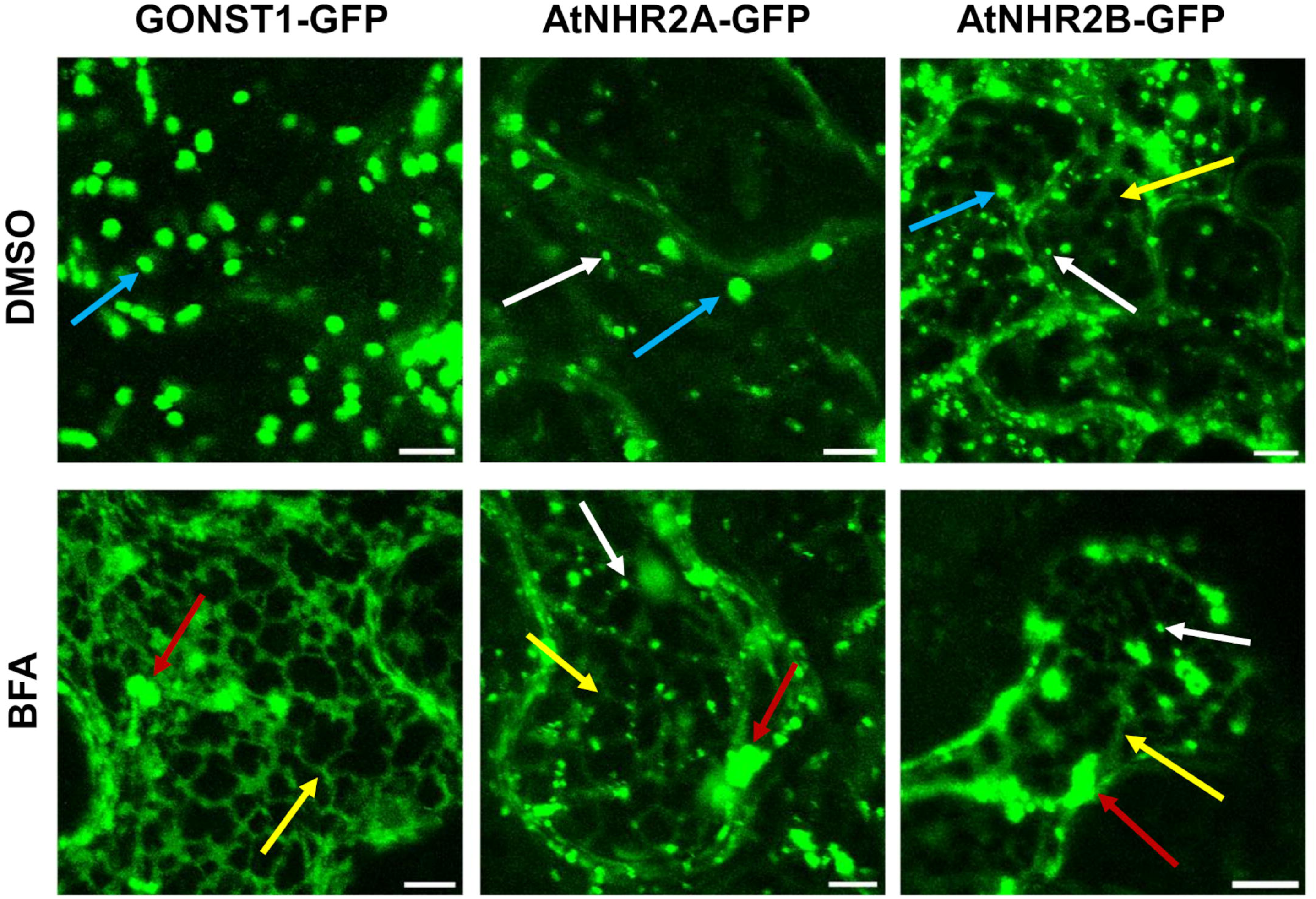
AtNHR2A and AtNHR2B proteins traffic through both Golgi-dependent and Golgi-independent pathways. *N. benthamiana* leaves transiently expressing *GONST1-GFP, AtNHR2A-GFP* and *AtNHR2B-GFP* for 2 days were treated with 0.1% DMSO (mock-treatment), or Brefeldin A (BFA) at 50µg/mL and imaged at 1h after BFA infiltration. Images show Golgi bodies (blue arrows), ER (yellow arrows), punctae (white arrows) and BFA compartments (red arrows). Images are Z-stacks of 7 single sections. Scale bar = 5µm.

Treatment with BFA retained *GONST1-GFP* signal in the net pattern of the ER (Figure 5, yellow arrow) and resulted in the formation of BFA compartments generated by the fusion of ER and Golgi stalks (Figure 5, red arrow). In the case of AtNHR2A-GFP and AtNHR2B-GFP, treatment with BFA generated the accumulation of the GFP signal in ER (Figure 5, yellow arrow), as well as the BFA compartment (Figure 5, red arrow), similar to the pattern of localization of *GONST1-GFP*, confirming that AtNHR2A-GFP and AtNHR2B-GFP move from ER to Golgi and that movement is prevented by BFA. Interestingly, in addition to the localization to ER and BFA compartment under BFA treatment, AtNHR2A-GFP and AtNHR2B-GFP signals were also found in punctae (Figure 5, white arrow), that was observed even after 3 h of BFA treatment (Supplementary Figure S1), indicating that these AtNHR2A-GFP- and AtNHR2B-GFP-containing punctae are BFA-insensitive and represent compartments that bypass Golgi. These results demonstrate that AtNHR2A and AtNHR2B transition through Golgi-dependent and Golgi-independent pathways, supporting their role in both CPS and UPS pathways, respectively.

### AtNHR2A and AtNHR2B traffic through the MVB-vacuole-plasma membrane route of the UPS pathway

The implication that both AtNHR2A and AtNHR2B participate in UPS pathway (Figures 2 and 5), led us to evaluate one possible UPS pathway involving MVB and vacuole. For that purpose, we co-expressed *AtNHR2A-GFP* and *AtNHR2B-GFP* with the MVB and vacuole membrane markers, *ARA6-mCherry* and Gamma tonoplast intrinsic protein *γTIP-mCherry*, respectively. Co-expression of *AtNHR2A-GFP* or *AtNHR2B-GFP* with the *ARA6-mCherry* revealed moderate levels of co-localization at MVB, as determined by PCC scores of 0.46±0.15 and 0.42±0.11, respectively, and tM1 and tM2 value in range of 0.6-0.7 (Figure 6A, Supplementary Table S5). Transient co-expression of *35S:AtNHR2A-GFP* or *35S:AtNHR2B-GFP* and *35S: ɣTIP-mCherry*, revealed strong co-localization as determined by high PCC values of 0.55±0.075 and 0.55±0.078, respectively, and tM1 and tM2 value between 0.7-0.9 (Figure 6B and Supplementary Table S5). The localization of these proteins to the tonoplast was confirmed by the visualization of AtNHR2A-GFP and AtNHR2B-GFP signals found at trans-vacuolar strands, observed as tubular structures that invaginate into vacuolar lumen (Madina et al., 2019) (Supplementary Figure S2).

**Figure 6.**
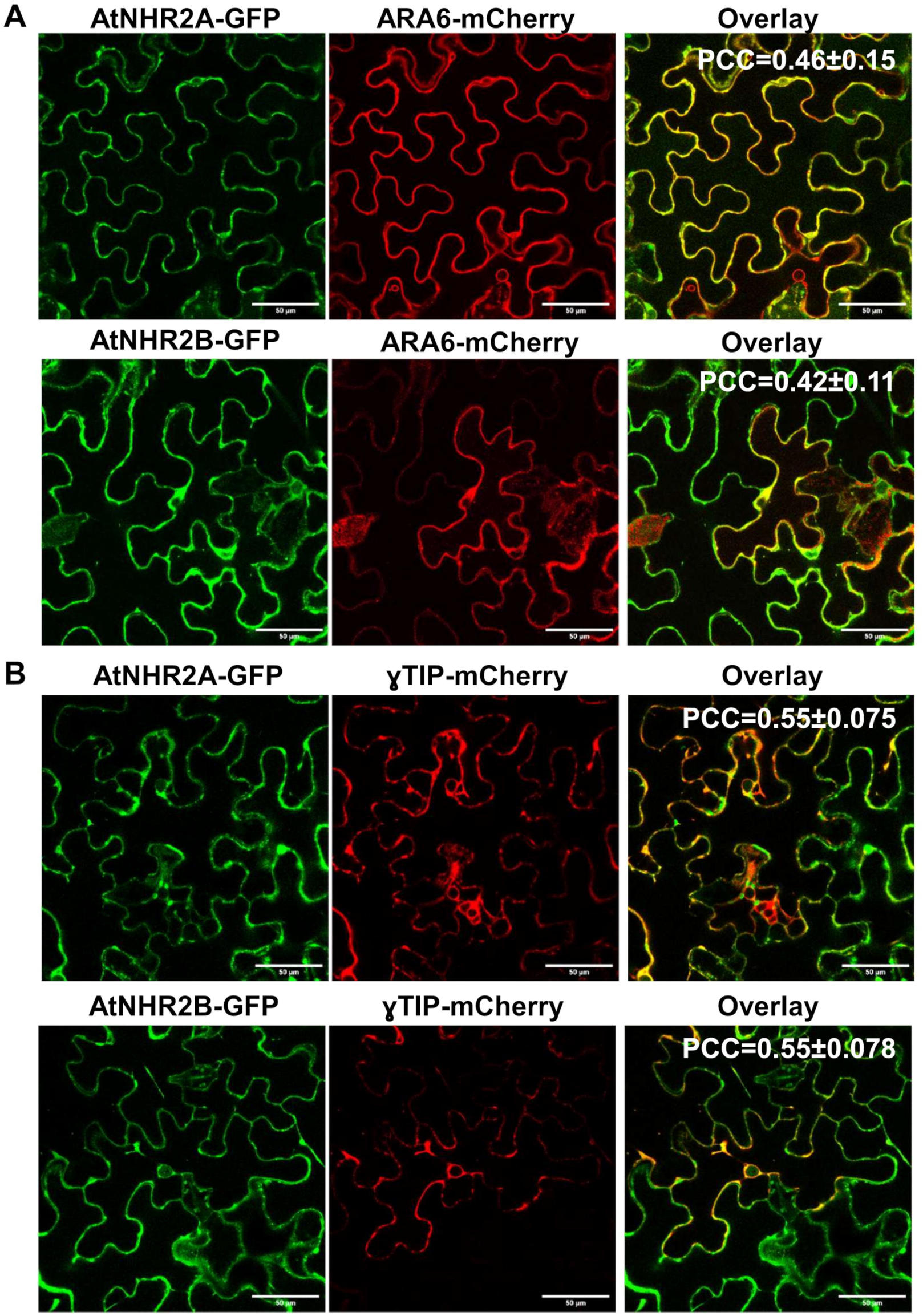
Co-localization of AtNHR2A and AtNHR2B with MVB and tonoplast markers. Transient co-expression in *Nicotiana benthamiana* was used to evaluate co-localization of AtNHR2A-GFP and AtNHR2B-GFP with the MVBs marker ARA6-mCherry (A) and Tonoplast marker ɣTIP -mCherry (B). The images were taken 3 days after infiltration in green and red channels using laser scanning confocal microscopy. Pictures are representative images. Pearson’s correlation coefficient (PCC) value represents the average of 10 microscopic fields. Scale bar = 50µm.

Our findings that AtNHR2A and AtNHR2B function in Golgi-independent pathways (Figure 2, 4, 5 and Supplementary Figure S1), along with the localization of AtNHR2A and AtNHR2B to MVB and tonoplast, led us to hypothesize that the trafficking pathway of AtNHR2A and AtNHR2B involves transition from MVBs to the central vacuole. To evaluate trafficking of AtNHR2A and AtNHR2B to the vacuole, we used the vesicle trafficking inhibitor Wortmannin (WM) that promotes the fusion of MVBs into enlarged MVBs; this homotypic fusion prevents the fusion between MVB and central vacuole interfering with the movement of proteins from MVB to vacuole. As a positive control, we used the MVB marker ARA6 that mediates transport from MVB to the vacuole (Ueda et al., 2004, Ebine et al., 2011). Transient expression of *ARA6-mCherry* or *AtNHR2A-GFP* or *AtNHR2B-GFP* and further treatment with 0.1% DMSO, showed fluorescent signal in punctae (Figure 7A, white arrows). WM treatment generated the accumulation of ARA6-mCherry signal into large spherical structures (Figure 7A, blue arrows), representing enlarged MVB originated from fusion of MVBs containing ARA6-mCherry (Figure 7A, blue arrows) as previously described (Ebine et al., 2011, Ueda et al., 2001). WM treatment also showed AtNHR2A-GFP and AtNHR2B-GFP signals accumulating in spherical structures similar with the ARA6-mCherry containing enlarged MVB (Figure 7A, blue arrows). Interestingly, WM treatment also caused accumulation of AtNHR2B-GFP in other larger spherical structures (Figure 7A, purple arrows), that could represent small vacuoles in route to fuse to the central vacuole but that were halted under WM treatment.

**Figure 7:**
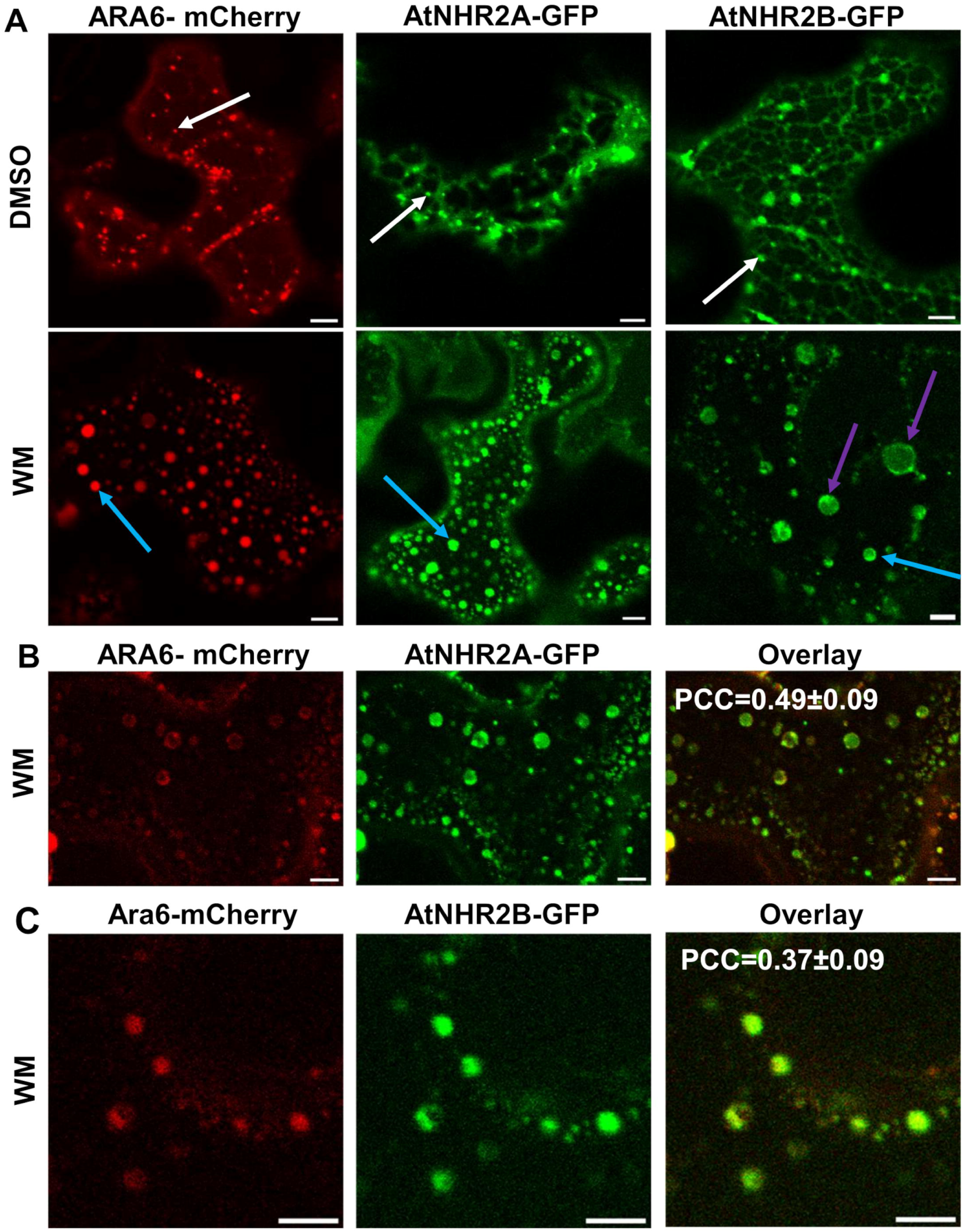
AtNHR2A-GFP and AtNHR2B-GFP are Wortmannin sensitive and Wm treatment triggers co-localization with MVB. *N. benthamiana* leaves transiently expressing *ARA6-mCherry, AtNHR2A-GFP* or *AtNHR2B-GFP* for 2 days were infiltrated with 0.1% of DMSO (mock-treatment) or 25 µM of Wortmannin (WM). mCherry and GFP signals were detected and imaged at 30 min post infiltration by using laser scanning confocal microscopy. White arrows represent punctae, blue arrows represent enlarged MVB, purple arrows represent small vacuoles. Images are single sections. Scale bar = 5µm (A). Transient co-expression of *Ara6-mCherry* with *AtNHR2A-GFP* (B) or *AtNHR2B-GFP* (C) followed by WM treatment was used to evaluate Wm-mediated co-localization. Pictures are representative images. Pearson’s correlation coefficient (PCC) value represents the average of 10 microscopic fields.

To confirm that the spherical structures harboring AtNHR2A-GFP and AtNHR2B-GFP signals observed after WM treatment were also enlarged MVB, we further co-expressed *ARA6-mCherry* with *AtNHR2A-GFP* (Figure 7B) or with *AtNHR2B-GFP* (Figure 7C) under WM treatment to evaluate their co-localization. The results showed co-localization of ARA6-mCherry and AtNHR2A-GFP with a PCC value between 0.49 ± 0.09 and tM1 and tM2 value of 0.6 which correspond to moderate levels of co-localization (Figure 7B and Supplementary Table S5). Similarly, co-localization between ARA6-mCherry and AtNHR2B-GFP resulted in the PCC value between 0.37 ±0.09 and tM1 and tM2 value as 0.45 and 0.4 (respectively) (Figure 7C and Supplementary Table S5), indicates partial colocalization. These results suggest a role for AtNHR2A and AtNHR2B transporting proteins to the vacuole via MVB, highlighting a specific route for UPS.

The last step in both CPS and UPS pathways involved fusion events with the plasma membrane. The implication that both AtNHR2A and AtNHR2B participate in both CPS and UPS pathways led us to evaluate their co-localization with the PM marker *AtPIP2A-mCherry.* We found that both *AtNHR2A-GFP* and *AtNHR2B-GFP* co-localize with *AtPIP2A-mCherry* at moderate levels as determined by PCC scores of 0.36±0.1 and 0.42±0.18, respectively, and tM1 and tM2 values of ∼0.7 corresponding to moderate co-localization (Figure 8, Supplementary Table S5). Collectively, our results demonstrate that AtNHR2A and AtNHR2B transition through endomembrane compartments to the vacuole and ultimately the plasma membrane, the convergent endpoint of CPS and UPS secretion.

**Figure 8.**
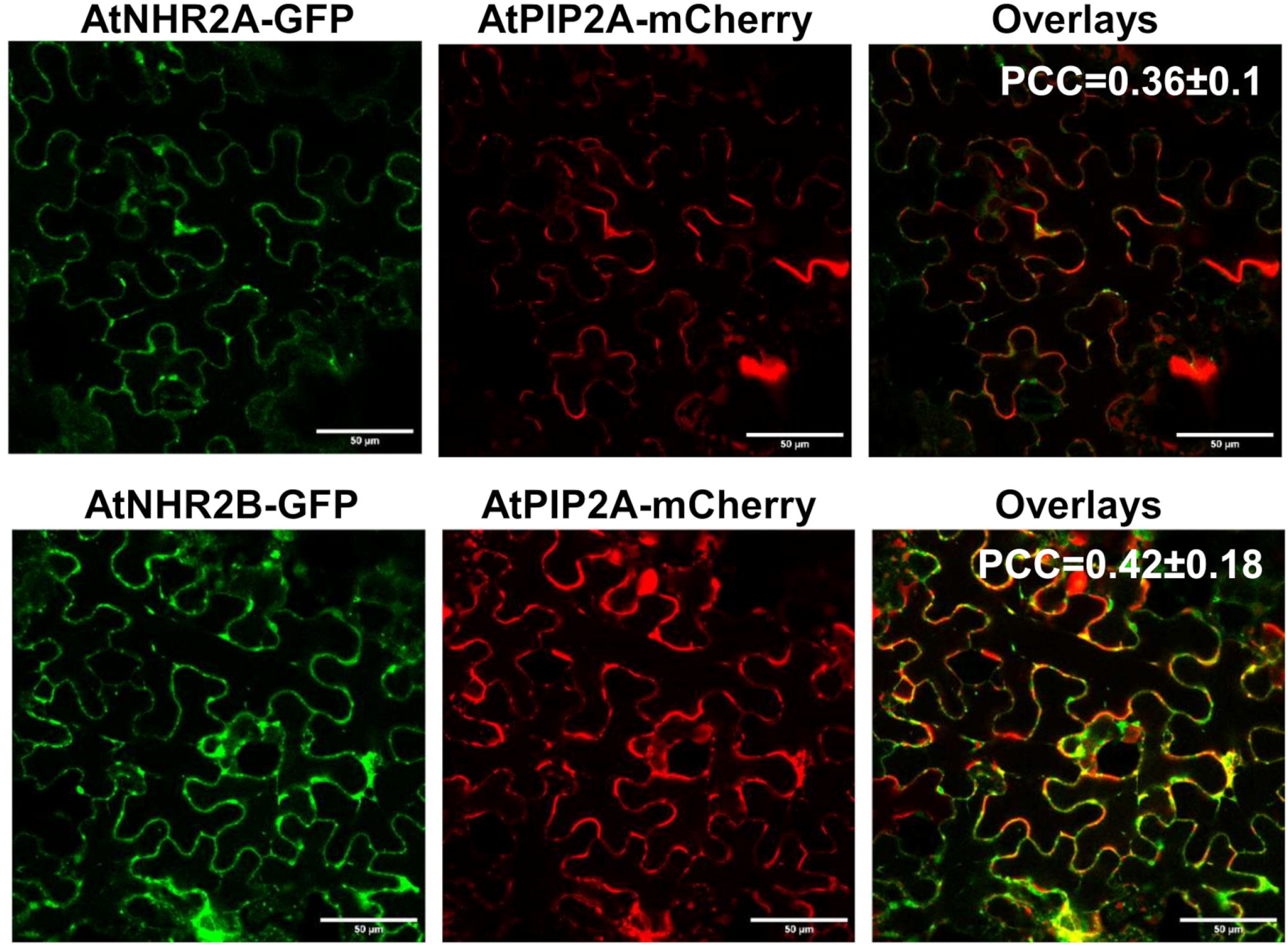
Co-localization of AtNHR2A and AtNHR2B with plasma membrane marker. Transient co-expression in *Nicotiana benthamiana* was used to evaluate co-localization of AtNHR2A-GFP and AtNHR2B-GFP with the plasma membrane marker AtPIP2A-mCherry. Infiltrated leaves were imaged at *2* days after infiltration in the green and red channels by laser scanning confocal microscopy. Pictures are representative images. Pearson’s correlation coefficient (PCC) value represents the average of 10 microscopic fields. Scale bar = 5µm.

## Discussion

Protein secretion to the apoplast is essential for the execution of critical defense responses such as cell wall fortification and release of antimicrobials, when plants are threatened by pathogens (Bhandari and Brandizzi, 2024, Gu et al., 2017, Kwon et al., 2008). The delivery of defense-related proteins to the apoplast occurs through CPS and UPS pathways that plants also employ for growth and development (Wang et al., 2018, Zhang et al., 2019, Kanazawa and Ueda, 2017). We previously showed that the immune-related proteins AtNHR2A and AtNHR2B participate in CPS to secrete subsets of defense-related proteins that not only contribute to cell wall reinforcement and antimicrobial activity but also restrict pathogens’ access to nutrients in the apoplast (Maia et al., 2026). In this study, we showed that AtNHR2A and AtNHR2B also participate in UPS, by identifying secreted proteins lacking the N-terminal secretion signal that were in lower abundance in the apoplast of the double mutant *Atnhr2bAtnhr2a* when compared with the apoplast proteome of wild type-Col-0. Similar to the AtNHR2A- and AtNHR2B-dependent apoplastic proteome associated with CPS that uncovered defense-related proteins, the AtNHR2A- and AtNHR2B-dependent apoplastic proteome associated with UPS also revealed three proteins associated with responses to biotic stress: Proteasome Regulator 1 (PTRE1) (Thulasi Devendrakumar et al., 2019), Actin depolymerization factor 2 (ADF2) (Clement et al., 2009), and S-formylglutathione hydrolase (SFGH) (Hütten et al., 2015). Strikingly, among the proteins secreted in AtNHR2A- and AtNHR2B-dependent manners through the UPS pathway, we uncovered eight proteins involved in abiotic stress responses: Responsive to ABA18 (RAB18) (Lång and Palva, 1992, Lang et al., 1994), Responsive to desiccation 29B (RD29B) (Yamaguchi-Shinozaki and Shinozaki, 1993, Msanne et al., 2011), Cold shock domain protein 1 (CSDP1) (Park et al., 2009, Juntawong et al., 2013), Homogentisate 1,2-dioxygenase (HGO) (Huang et al., 2018, Han et al., 2013), UDP-glycosyltransferase 71B7 (UGT71B7) (Dong et al., 2014), GA-stimulated Arabidopsis 4 (GASA4) (Ko et al., 2007, Rubinovich and Weiss, 2010), TOM1-like protein 3 (TOL3) (Korbei et al., 2013, Moulinier-Anzola et al., 2014), and Probable glutathione peroxidase 8 (GPX8) (Gaber et al., 2012). Interestingly, among those proteins we found CSDP1 previously identified as an AtNHR2A interactor (Singh et al., 2020), suggesting that CSDP1 might be a putative cargo transported in AtNHR2A-vesicles. Two additional proteins that have been implicated in both abiotic and biotic stress responses include: Peptidyl-prolyl cis-trans isomerase (ROC3) (Liu et al., 2021, Pogorelko et al., 2014) and BPA1-like 1 (PBL1) (Li et al., 2019, Palm et al., 2019). Intriguingly, proteins associated with biotic stress responses were only found at 24 hai. At 72 hai, we found two proteins associated with plant responses to salt stress: DEK domain containing protein 3 (DEK3) (Waidmann et al., 2014, Waidmann et al., 2022) and Protein Kinasse (SnRK2.4) (Mazur et al., 2021, McLoughlin et al., 2012) and another protein (NAD(P)-binding Rossmann-fold superfamily protein (CCR-like 1) implicated in lignin biosynthesis (Raes et al., 2003), which could be important in cell wall fortification during defense responses. Our findings that AtNHR2A and AtNHR2B are required for secretion of proteins associated with plant responses to biotic and abiotic stresses, in addition to the identification of seven proteins that operate through the stress hormone ABA, suggests that AtNHR2A and AtNHR2B may act as a convergence point coordinating secretion in response to multiple environmental cues through ABA (Bharath et al., 2021).

Transport of cargo through subcellular compartments involves cargo selection in vesicles formed by budding from a donor compartment, vesicle movement to a cellular destination, vesicle tethering to target membranes at destination and vesicle fusion to target membranes (Bonifacino and Glick, 2004). Since we did not find AtNHR2A or AtNHR2B in the apoplast, it is unlikely that these proteins are vesicles cargo. Instead, the presence of transmembrane domains in these proteins (Singh and Rojas, 2018), together with our finding herein that AtNHR2A and AtNHR2B transition through multiple endomembrane compartments, suggest that these proteins accompany their cargoes through endomembrane compartments possibly to mediate membrane-related processes such as vesicle budding, tethering or fusion to target membrane.

Dissecting the specific trafficking routes for AtNHR2A and AtNHR2B, we found that these proteins transition through successive endomembrane compartments including ER, Golgi, MVBs and vacuole. Since MVBs and vacuole are not part of CPS (Drakakaki and Dandekar, 2013), our findings highlight that the UPS routes for AtNHR2A and AtNHR2B are either MVB-vacuole-PM and/or MVB-PM. While our current data does not allow us to discriminate between those options, it strengthens the role of MVB and vacuole in plant immunity in processes such as cell wall reinforcement and release of proteins with antimicrobial properties (An et al., 2006, Hatsugai et al., 2009). Our finding that pathogen inoculation resulted in the secretion of proteins previously implicated in plant responses to abiotic stress is puzzling but highlights the potential role of AtNHR2A and AtNHR2B contributing to plant adaptation under varying environmental conditions.

The strong co-localization of AtNHR2A and AtNHR2B with the MVB marker Ara6 is particularly interesting, because just like Ara6, AtNHR2A and AtNHR2B are plant specific proteins (Ebine et al., 2011, Ueda et al., 2001), suggesting that these three proteins could function together in biological processes unique to plants. Ara6 is a Rab GTPase that, similar to other Rab GTPases functioning in endomembrane trafficking, control the specificity and direction of vesicle transport (Minamino and Ueda, 2019, Zhen and Stenmark, 2015) with Ara6 specifically mediating trafficking from MVB to plasma membrane and vacuole (Bottanelli et al., 2011, Bottanelli et al., 2012, Ebine et al., 2011). Ara6 function was previously associated with plant responses to salt and osmotic stress (Ebine et al., 2012, Ebine et al., 2011, Tsutsui et al., 2015), that could be linked to our identification of subsets of apoplastic secreted proteins such as RD29B, CSDP1, HGO, UGT71B7 and PBL1 known to participate in these responses (Msanne et al., 2011, Park et al., 2009, Palm et al., 2019, Dong et al., 2014, Huang et al., 2018). In the context of plant responses to biotic stress, overexpression of Ara6 resulted in enhanced callose deposition that effectively restricted proliferation of the fungal pathogen *Golovinomyces orontii* (Inada et al., 2016, Inada et al., 2017), and we also previously found that AtNHR2A and AtNHR2B play a role in callose deposition in response to non-adapted bacterial pathogens (Singh et al., 2018). Thus, although the specific functions of AtNHR2A and AtNHR2B are still not known, it is possible these functions could be linked to the RAB GTPase function of Ara6 tethering vesicles to target membranes (Ebine et al., 2012). Rab GTPases cycle between a cytosolic GDP (<u>g</u>uanosine <u>dip</u>hosphate)-bound (inactive) state to a membrane-bound GTP (<u>g</u>uanosine tri<u>p</u>hosphate)-bound (active) state (Stenmark, 2009), and in this membrane-bound state interact with various proteins known as Rab effectors that control the specificity of the transport to a given subcellular compartment (Zhen and Stenmark, 2015). We speculate that AtNHR2A and AtNHR2B could be an Ara6 Rab effector, more research is needed to confirm this.

Altogether, this study revealed that AtNHR2A and AtNHR2B participate in Golgi-dependent and Golgi-independent trafficking pathways associated with CPS and UPS and uncovered their participation in MVB-vacuole-mediated UPS necessary for the secretion of proteins responsible for plant responses to abiotic and biotic stresses.

## Acknowledgements

We would like to thank to Dr. Jaideep Mathur for providing *mEOS-GONST* construct, Dr. Elison Blancaflor for providing *ARA6-mCherry* construct, Dr. Bara Altartouri at the Nebraska Center for Biotechnology, Microscopy Core Facility and Dr. Kwang-moon Cho with assistance with imaging experiments.

## Author contributions

CR, HTKN,TM and BO, designed the research; HTKN, TM and BO, performed the experiments and analyzed the data; CR, HTKN,TM and BO analyzed and interpreted the data; CR, HTKN,TM and BO, wrote and revised the manuscript.

## Funding

This work was supported by the National Science Foundation CAREER award number 2332080 and the University of Nebraska-Lincoln start up support to CR.

## Conflict of interest

None declared

## Data availability

The authors declare that all the data involved have been provided in the main text or in the Suppporting Information.

## Notes

### Competing Interest Statement

The authors have declared no competing interest.

