## Supplementary figures and images for "AtNHR2A and AtNHR2B participate in unconventional protein secretion in response to environmental stress"

### Supplemental Figure 1

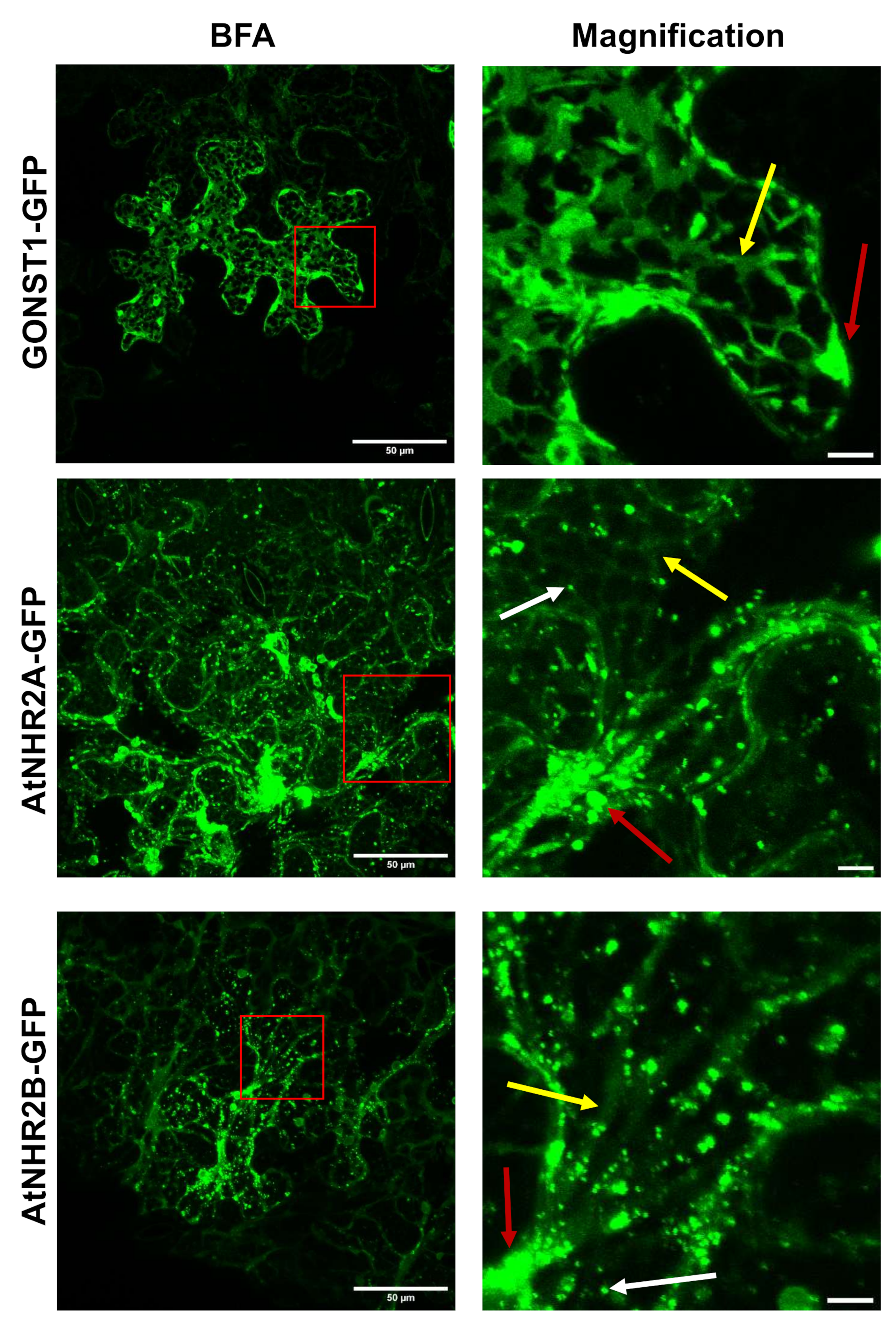

### Supplemental Figure 2

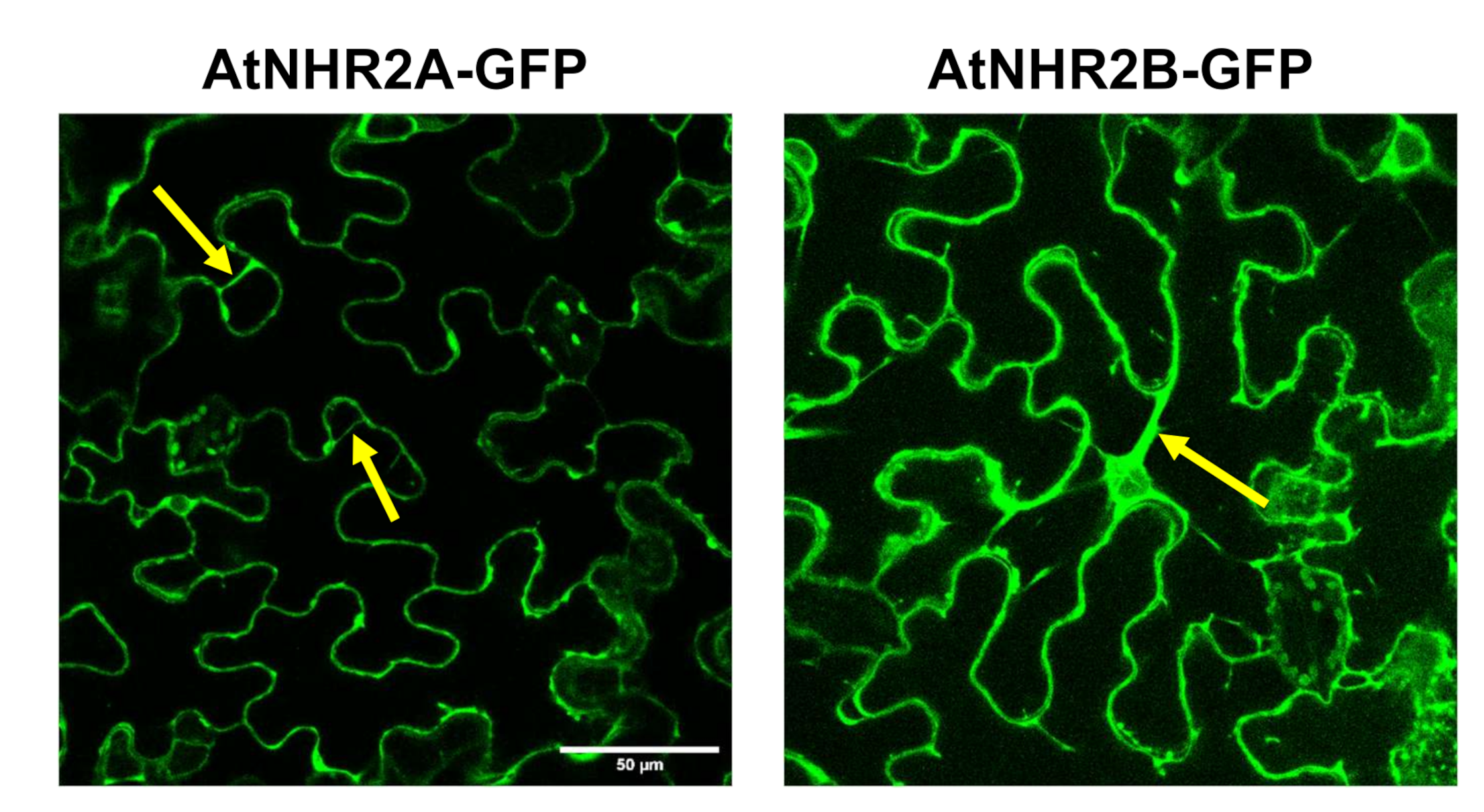
